# Divergent sperm traits in Carabidae ground beetles (Insecta: Coleoptera)

**DOI:** 10.1101/817635

**Authors:** Kôji Sasakawa

## Abstract

Sperm exhibit marked morphological diversity, and investigations into sperm diversity can further the understanding of many areas of evolutionary biology. In this study, using light microscopy, sperm morphology was examined in 39 species of Carabidae from eight subfamilies, including five subfamilies in which sperm morphology has not previously been examined. In all but one of the subfamilies, the subfamily members shared the same type of sperm: single sperm were observed in Cicindelinae, Nebriinae, and Trechinae; sperm conjugates, in which numerous sperm adhere together, were observed in Elaphrinae, Patrobinae, and Brachinae; and both single sperm and sperm conjugates were observed in Broscinae. In the remaining subfamily, Harpalinae, most species formed sperm conjugates, but some species formed single sperm. Some noteworthy sperm were also observed: the shortest single sperm in the order Coleoptera was found; multiflagellated sperm were observed, which had previously been reported from only one species in the class Insecta; and size variation of sperm conjugates, which may represent size dimorphism, was observed. Based on the results of this and previous studies, the evolutionary pattern of sperm traits and the phylogenetic utility of sperm morphologies in Carabidae are discussed.

## INTRODUCTION

Insects possess divergent reproductive traits, which are of interest in many areas of evolutionary biology. In these reproductive traits, male sperm show marked morphological diversity, and this diversity is associated with two research topics. The first addresses the evolution of reproductive traits. Reproductive traits often exhibit correlated evolution, and sperm also demonstrates morphological association with other reproductive traits (Simmons, 2001; Pitnick *et al*., 2009). Thus, the study of sperm morphology is necessary to develop an understanding of the mechanisms by which diversity in reproductive traits occurred. The second research topic involves phylogenetic analyses. For example, among higher taxa, it is often difficult to determine the homology of morphological characters, which hampers phylogenetic reconstruction based on morphological data. However, even in such cases, the homology of sperm morphology can often be determined, and comparative morphology can provide insights into the reconstruction of phylogenies (i.e., spermiocladistics) (Jamieson *et al*., 1999). Nevertheless, despite the utility of these studies, sperm morphology remains unexamined in many insect groups.

The beetle family Carabidae is one such group. It comprises more than 40,000 species that are classified into about 20 subfamilies according to current taxonomic systems; members of the group inhabit a variety of habitats and exhibit a diverse range of morphologies and life histories (Lövei & Sunderland, 1993; Löbl & Löbl, 2017). Despite this diversity, sperm morphology has been examined in only four subfamilies (Cicindelinae, Scaritinae, Carabinae, and Harpalinae) (Sasakawa & Toki, 2008; Sasakawa, 2009a). In addition, sufficient numbers of species for within-group comparative studies have only been examined for two groups, the Carabinae tribe Carabini and the Harpalinae tribe Pterostichini; only one or a few species have been examined in other taxa. However, the results of previous studies indicate the importance of studying sperm morphology in Carabidae. For example, in the Carabini species studied (genus *Carabus* subgenus *Ohomopterus*), the size of a male genital organ, the “copulatory piece,” is associated with the size and dimorphism of the sperm bundle, which is a type of conjugated sperm (Takami & Sota, 2007). In the Pterostichini, the size of the female spermatheca is positively correlated with the size of the sperm bundle (Sasakawa, 2007). These reports suggest correlated evolution between the sperm and genitalia, and indicate that examination of sperm morphology is necessary in studies investigating the evolution of reproductive traits in Carabidae. Studies of sperm morphology are also important for elucidation of the within-family phylogenies of Carabidae. Sasakawa & Toki (2008) compared sperm morphology among species from five tribes of three subfamilies, whose phylogenetic positions within the family were elucidated by molecular phylogenetic analysis; their results revealed that conjugated sperm with an elongated central rod may be an autapomorphy of Harpalinae, the largest and most derivative subfamily of Carabidae (Löbl & Löbl, 2017). This assumption was supported by subsequent studies of a species from an additional tribe within Harpalinae (Sasakawa, 2009b), and a species from an additional subfamily from basal lineages (Sasakawa, 2009a). These reports imply the utility of sperm morphology for reconstructing phylogenies.

In this study, using light microscopy sperm morphology was examined in 39 carabid species of eight subfamilies, including five subfamilies for which sperm morphology has not been examined. Although the numbers of species examined in each group were limited, the results of this and previous studies encompass many major lineages of Carabidae. Thus, the results provide insight into the evolutionary patterns and phylogenetic utility of sperm morphology, as well as enabling further detailed studies of each group.

## MATERIALS AND METHODS

Three species of Nebriinae, one species of Cicindelinae, two species of Elaphriinae, one species of Broscinae, four species of Trechinae, two species of Patrobinae, one species of Brachinae, and twenty-five species of Harpalinae were examined (Table 1). Live reproductive adult males were dissected within 2 days after field collection; the seminal vesicles were removed from the reproductive organs, and sperm obtained from the seminal vesicles were examined. Separation of the seminal vesicles from the body was performed in a Petri dish, and dissection of the seminal vesicles and subsequent observation of live sperm were performed on a glass slide, both in Ringer’s solution using sharp tweezers. Live sperm were observed and photographed with a light microscope, followed by Giemsa staining for a detailed morphological investigation. The sperm obtained from the seminal vesicles were considered to reflect the characteristics of the sperm within the ejaculate, such as the spermatophore, considering the results of previous studies in other species of Carabidae, in which no morphological differences were found between sperm obtained surgically from the seminal vesicles and that found in the ejaculate (Takami, 2002; Sasakawa, 2009a).

**Table 1.**
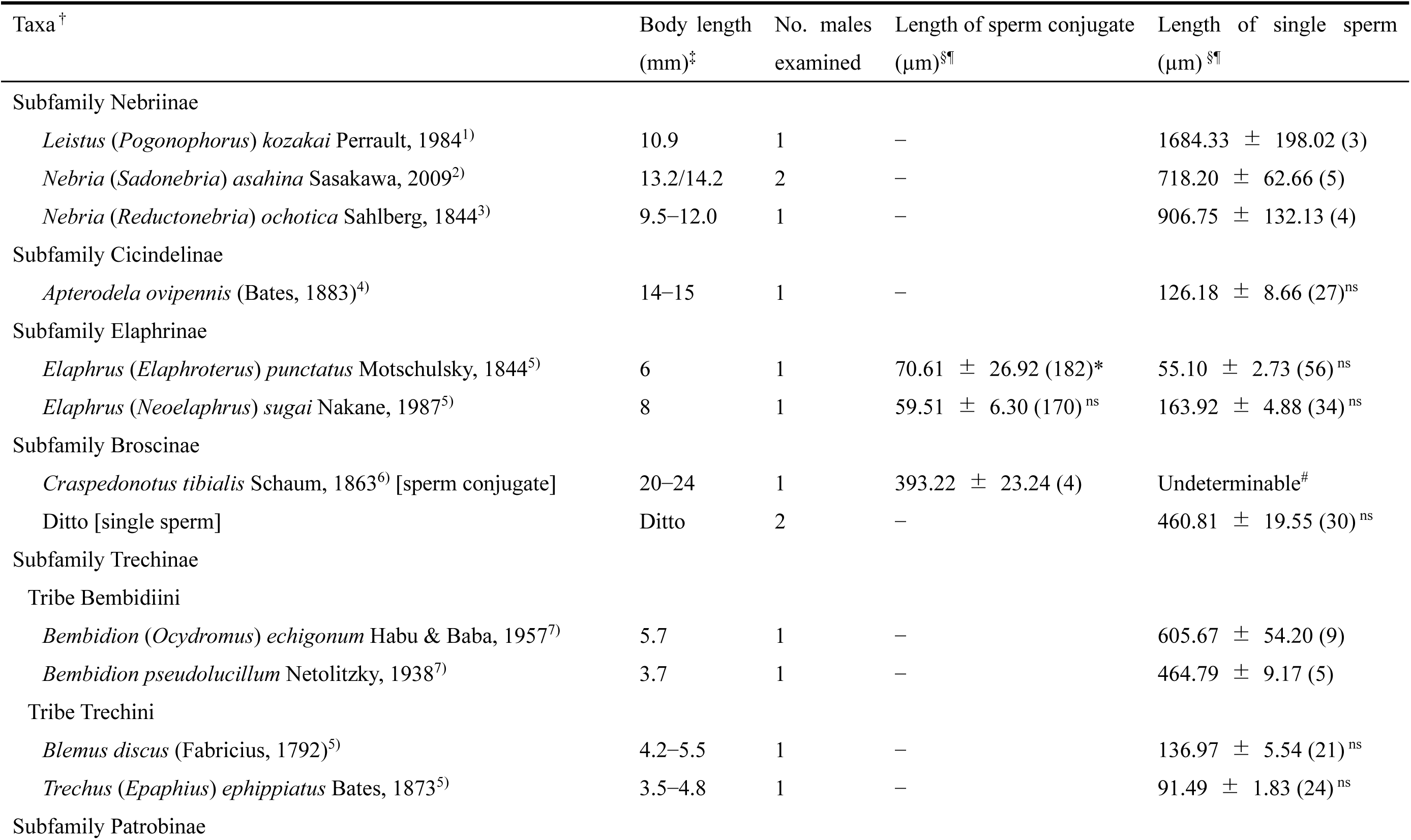
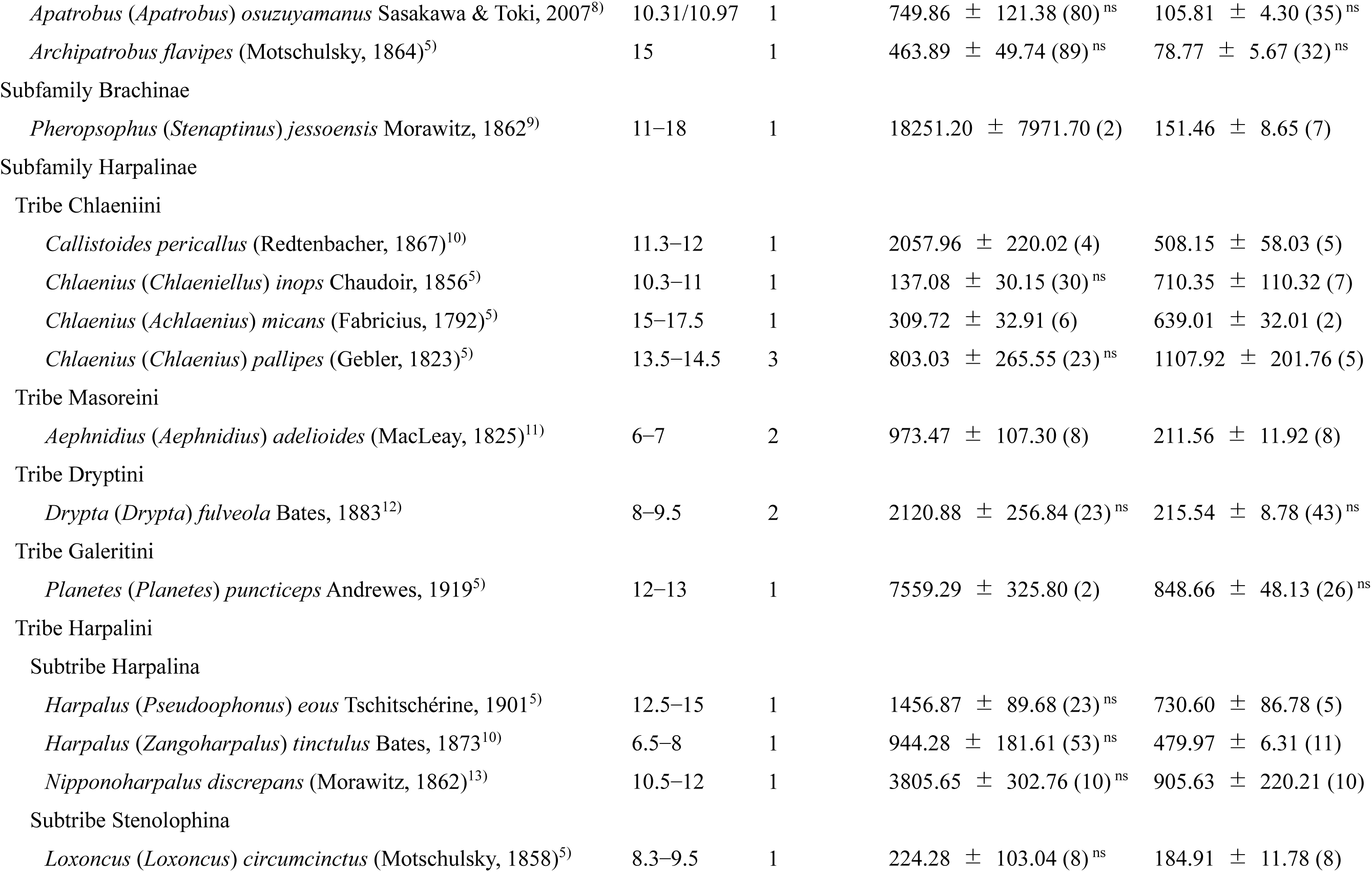
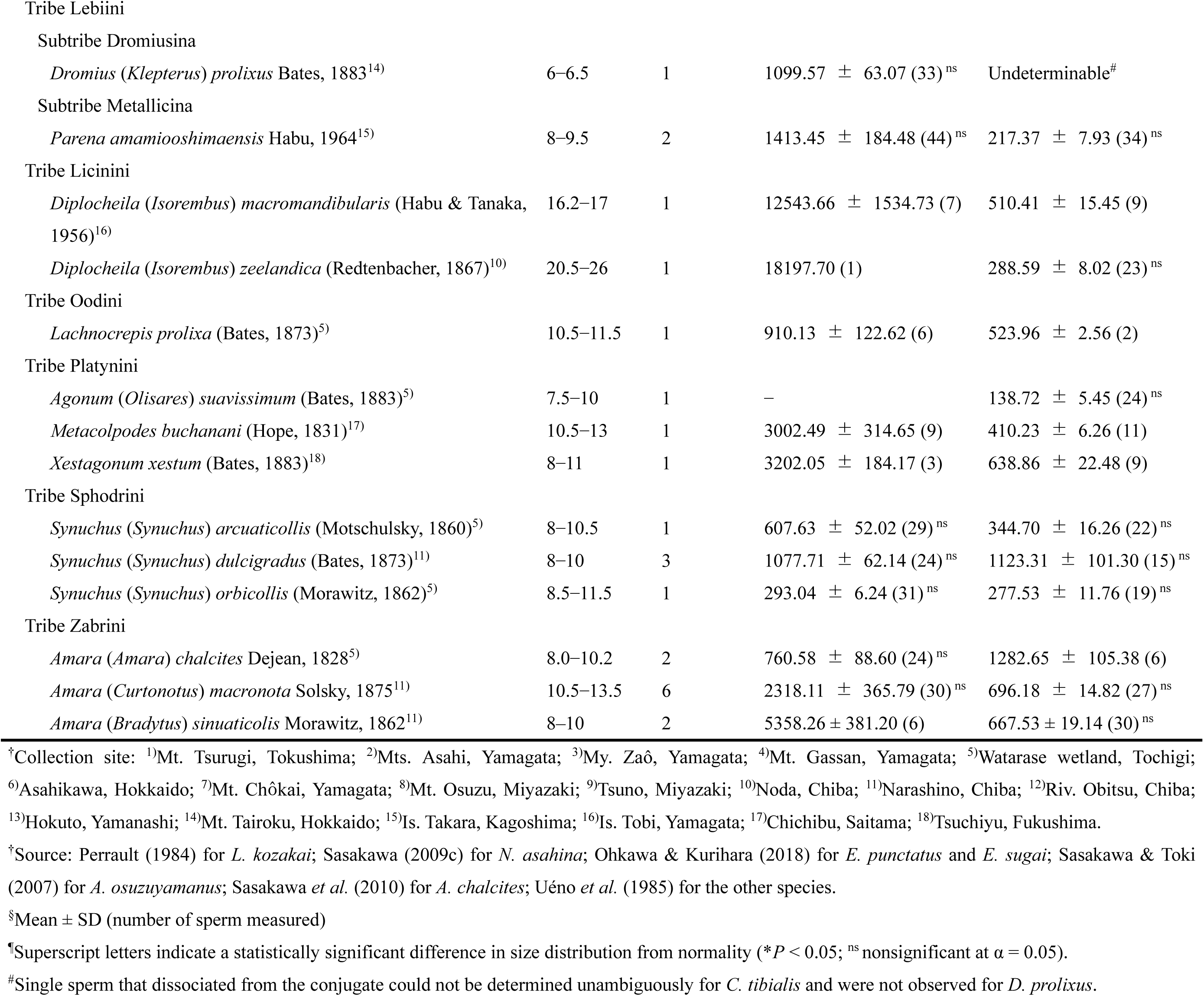
Species data

To describe the morphology, two terms are defined herein. First, all conjugated sperm are referred to as “sperm conjugate”. Higginson & Pitnick (2011) noted that many different, often taxon-specific terms have been used to refer to conjugated sperm and, for the avoidance of confusion, proposed the use of two categories, primary conjugate and secondary conjugate, both of which include several subcategories, based on their developmental mechanisms. Because it is not yet clear whether the species studied here fall into the primary or secondary conjugate categories, this report uses the broad category that includes both types of conjugate, which is also used in Higginson & Pitnick (2011). Additionally, the structure to which individual sperm adhere in the sperm conjugate is referred to in this report as “spermatostyle”. Although various terms have been used to describe this structure, mainly depending on the shape (e.g., rod or cap), a single term is used here, irrespective of the shape. In some species, the structure that appears to correspond to the spermatostyle was observed, but adhesion of individual sperm on the surface was not. Here, those structures are also referred to as spermatostyle for convenience.

Length measurements were performed on Giemsa-stained specimens for both single sperm and sperm conjugate. Typically, even species that form sperm conjugate have a small number of single sperm in the seminal vesicle, as reported in Dytiscidae beetles (Higginson *et al*., 2012). In addition, on Giemsa-stained specimens of sperm conjugate, single sperm that dissociated from the spermatostyle during the staining procedure were often observed. These sperm were used for the measurement of single sperm, in species with sperm conjugate. For multiflagellated sperm that were found in a species of Platynini, the measurement was taken as the anterior end of the sperm head to the posterior end of the longest flagellum. In all but two species, the sperm conjugate size was measured from the anterior end to the posterior end of the spermatostyle; in the exceptions (*Elaphrus punctatus* and *Craspedonotus tibialis*), due to the indistinct border between the spermatostyle posterior end and the attached sperm, the posterior end used for measurement was newly established on the rearmost attached sperm (the posterior end of the sperm head for *E. punctatus*, and the tail end for *C. tibialis*). Spermatostyle length was also measured for species in which sperm were not attached to the spermatostyle. Because the sample size from single individuals was small in most cases, data from multiple individuals were pooled by species, assuming homogeneity of data among individuals. For single sperm and sperm conjugate with a sample size ≥ 15, the size distribution was assessed by testing goodness-of-fit to a normal distribution using the Kolmogorov-Smirnov test, with subsequent fitting of smooth curves to the histograms. ImageJ software (version 1.50i) (Rasband, 2016) was used to obtain the measurements, and R software (version 3.2.1) (R Development Core Team, 2015) was used for analysis thereof. Body length information was obtained from published literatures.

## RESULTS

### DESCRIPTIONS OF SPERM MORPHOLOGY

#### SUBFAMILY NEBRIINAE

(Fig. 1A, B)

**Figure 1.**
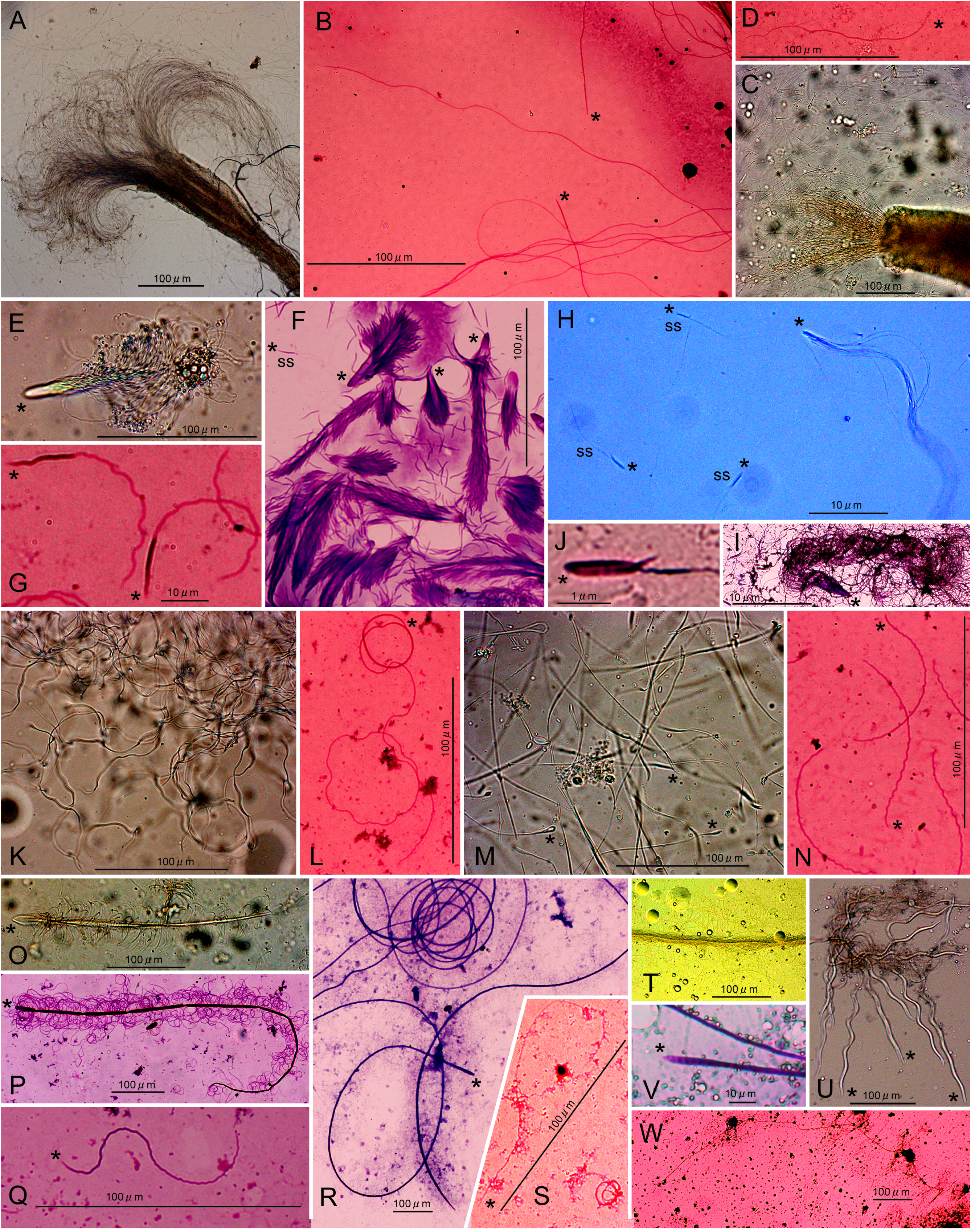
Sperm of various species of Carabidae observed in Ringer’s solution (RS) or with Giemsa stain (GS): *Leistus kozakai* single sperm in RS (A); *Nebria asahina* single sperm with GS (B); *Apterodela ovipennis* single sperm in RS (C) and with GS (D); an *Elaphrus sugai* sperm conjugate in RS (E); *Elaphrus punctatus* sperm conjugate (F) and single sperm (G) with GS; *Craspedonotus tibialis* sperm conjugate and single sperm in RS (H), and sperm conjugate (I) and single sperm (J) with GS; *Bembidion echigonum* single sperm in RS (K); a *Bembidion pseudolucillum* single sperm with GS (L); *Blemus discus* single sperm in RS (M); *Trechus ephippiatus* single sperm with GS (N); an *Archipatrobus flavipes* sperm conjugate in RS (O); *Apatrobus osuzuyamanu*s sperm conjugate (P) and single sperm (Q) with GS; *Pheropsophus jessoensis* sperm conjugate (R) and single sperm (S) with GS; a *Callistoides pericallus* sperm conjugate in RS (T); *Chlaenius micans* single sperm and spermatostyles in RS (U); *Chlaenius pallipes* spermatostyle (V) and single sperm (W) with GS. The heads of the single sperm and sperm conjugate, and the anterior ends of the spermatostyles without attached sperm, are indicated by an asterisk. The “ss” in the photos including sperm conjugate indicates single sperm that co-occur (for *C. tibialis*) or single sperm that dissociated from sperm conjugate (for the other species).

In all three species, only motile single sperm were observed (Fig. 1A, B). The heads of the sperm of the two *Nebria* species were slightly broader than the other parts of the sperm (Fig. 1B), but in the *Leistus* species, the head and other parts did not differ in width.

#### SUBFAMILY CICINDINAE

(Fig. 1C, D)

Only single sperm were observed (Fig. 1C). The sperm were motile and had heads that were only slightly broader than the tails (Fig. 1D).

#### SUBFAMILY ELAPHRINAE

(Fig. 1E–G)

In both species, sperm conjugate formed, in which the heads of numerous sperm were attached to a spermatostyle, while the tail of each sperm was free-moving (Figs. 1E, F), and single sperm that dissociated from the conjugate had an obviously broad head (Fig. 1G).

#### SUBFAMILY BROSCINAE

(Fig. 1H–J)

Both sperm conjugate and single sperm were observed (Fig. 1H). In sperm conjugate, the heads of numerous sperm were glued together, while the tail of each sperm was free-moving (Fig. 1I). Unlike the free single sperm described below, conjugate-forming single sperm did not have a differentiated head. Single sperm that co-occurred with the sperm conjugate had an elongated drop-shaped head, and a motile tail was attached to the head at a position in front of the posterior end of the head (Fig. 1J). On Giemsa-stained specimens, single sperm detached from the conjugate and free single sperm that lost their heads during specimen preparation were virtually indistinguishable.

#### SUBFAMILY TRECHINAE

(Fig. 1K–N)

Only motile single sperm were observed in both Bembidiini (Fig. 1K, L) and Trechini (Fig. 1M, N). The heads of the Bembidiini sperm were slightly broader than the other parts (Fig. 1L), whereas the heads of the Trechini sperm did not broaden (Fig. 1N).

#### SUBFAMILY PATROBINAE

(Fig. 1O–Q)

Sperm conjugate was observed, in which the heads of numerous sperm were attached to an elongated spermatostyle, while the tail of each sperm was free-moving (Fig. 1O, P). The spermatostyle was twisted in a right-handed direction at regular intervals, but was flexible and showed no conspicuous spiral structure. The head of the spermatostyle was spoon-shaped. The head of the detached single sperm was slightly broader than the other parts (Fig. 1Q).

#### SUBFAMILY BRACHINAE

(Fig. 1R, S)

Sperm conjugate was observed, in which the heads of numerous sperm were attached to a spermatostyle, while the tail of each sperm was free-moving (Fig. 1R). The spermatostyle was markedly elongated, without a conspicuous spiral structure. The head of the spermatostyle had a match-head shape. The head of the single sperm that dissociated from the conjugate was slightly broader than the other parts (Fig. 1S).

#### SUBFAMILY HARPALINAE

##### TRIBE CHLAENIINI

(Fig. 1T–W)

In *Callistoides pericallus*, sperm conjugate was observed, in which the heads of numerous sperm were attached to the spermatostyle, while the tail of each sperm was free-moving (Fig. 1T). The spermatostyle was a straight, rigid rod, and its head had a match-head shape. In the other three species, although spermatostyles were observed, only single sperm were present, which were unattached to the spermatostyles. These three species shared a feature not found in the other species examined: single sperm were longer than the spermatostyle. However, the shape of the spermatostyle differed between the species, with a wave form in *Chlaenius micans* (Fig. 1U), a short rod in *Chlaenius inops*, and a long rod in *Chlaenius pallipes* (Fig. 1V). The head of single sperm did not broaden (Fig. 1W).

##### TRIBE MASOREINI

(Fig. 2A–C)

**Figure 2.**
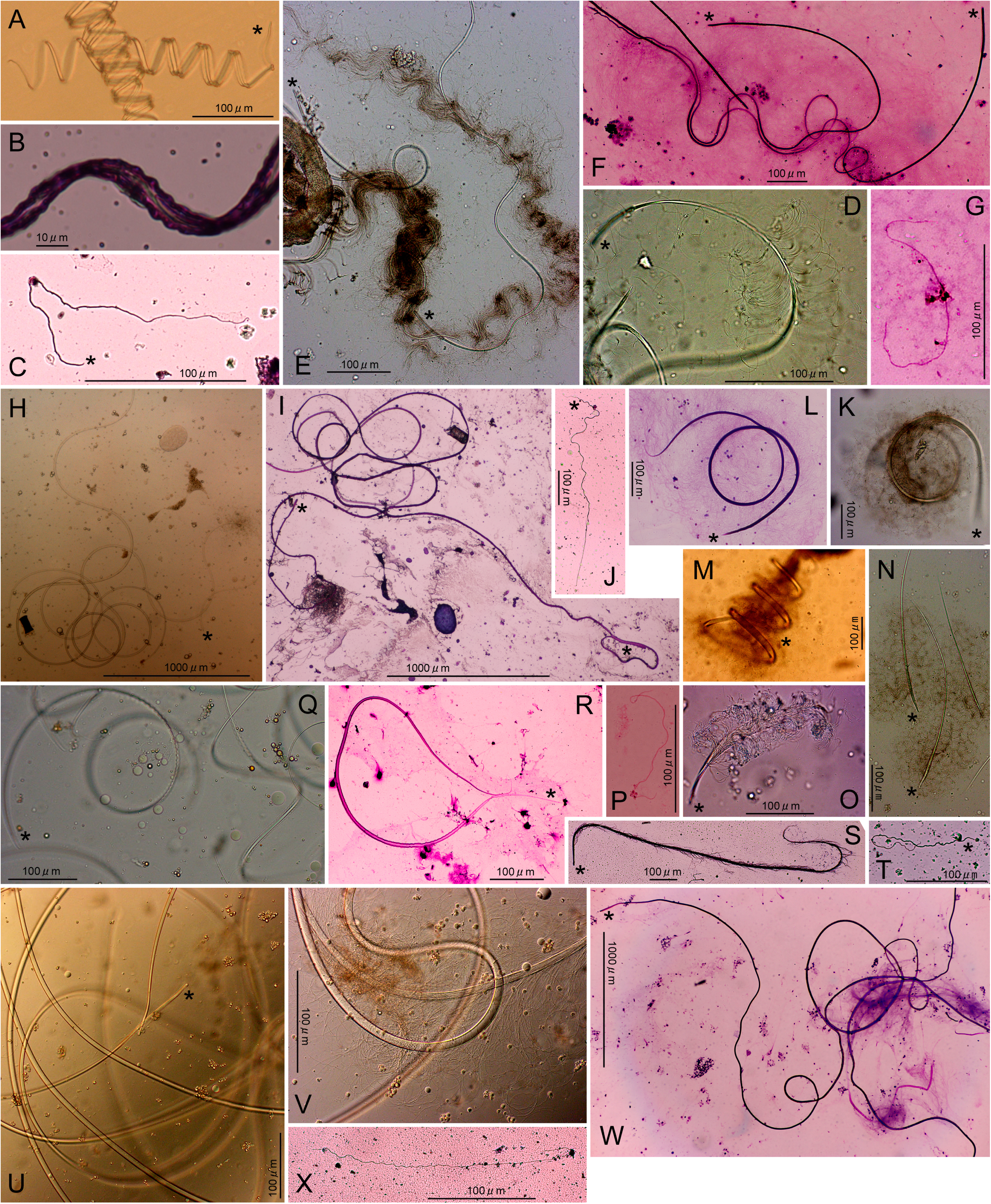
Sperm of various species of Carabidae: *Aephnidius adelioides* sperm conjugate in RS (A) and with GS (B), and single sperm with GS (C); *Drypta fulveola* spermatostyle with (D) and without (E, F) sperm on the surface, and single sperm (G) observed in RS (D, E) and with GS (F, G); *Planetes puncticeps* spermatostyles and single sperm in RS (H) and with GS (I), and single sperm with GS (J); *Harpalus eous* sperm conjugate in RS (K) and with GS (L); a *Nipponoharpalus discrepans* sperm conjugate in RS (M); *Harpalus tinctulus* sperm conjugate in RS (N); *Loxoncus circumcinctus* sperm conjugate in RS (O) and single sperm with GS (P); *Dromius prolixus* sperm conjugate in RS (Q) and with GS (R); *Parena amamiooshimaensis* sperm conjugate (S) and single sperm (T) with GS; *Diplocheila macromandibularis* spermatostyle with (U) and without (V) sperm on the surface in RS, and spermatostyles and single sperm with GS (W); *Diplocheila zeelandica* single sperm with GS (X). Explanations of the markings are the same as in Figure 1.

Sperm conjugate with a right-handed helical spermatostyle was observed (Fig. 2A). The sperm formed a sheet-like structure with a wavy edge, and adhered to both lateral sides of the spermatostyle (Fig. 2B). The head of the spermatostyle narrowed at the anterior end. The head of the detached single sperm was slightly broader than the other parts (Fig. 2C).

##### TRIBE DRYPTINI

(Fig. 2D–G)

Although some sperm were attached to the spermatostyle by their head (Fig. 2D), while their tail was free-moving, most sperm did not attach to the spermatostyle (Fig. 2E, F). The spermatostyle was elongated, without a conspicuous spiral structure. The spermatostyle head was slightly broader than the other parts. The heads of the single sperm did not broaden, and thus it was unclear which end was the head (Fig. 2G).

##### TRIBE GALERITINI

(Fig. 2H–J)

Although a spermatostyle was observed, the single sperm that were present were not attached to the spermatostyle surface. The spermatostyle was elongated, without a conspicuous spiral structure, and its head was slightly broader than the other parts (Fig. 2H, I). The head of single sperm was slightly broader than the other parts (Fig. 2J).

##### TRIBE HARPALINI

(Fig. 2K–P)

Sperm conjugate was observed in all species. The spermatostyle had a left-handed helical shape (Fig. 2K–M) or a rod shape (Fig. 2N, O). In both types, the head of the spermatostyle did not broaden, and numerous sperm were attached by their heads to the spermatostyle surface, while the tails were free-moving. The heads of single sperm that dissociated from the spermatostyle did not broaden, and thus it was unclear which end was the head (Fig. 2P).

##### TRIBE LEBIINI

(Fig. 2Q–T)

In both species, sperm conjugate with an elongated spermatostyle was observed, and the head of the spermatostyle narrowed at the anterior end. In the Dromiusina, the spermatostyle had a right-handed helical shape, and the sperm formed a sheet-like structure with a wavy edge and adhered to both lateral sides of the spermatostyle (Fig. 2Q, R). The sheet-like structures had firm adhesion, and single sperm that dissociated from the spermatostyle were not observed in the specimens examined. In the Metallicina, the spermatostyle was flexible, without a conspicuous spiral structure, and numerous sperm were attached by their heads to the spermatostyle surface, while the tails were free-moving (Fig. 2S). The head of the single sperm detached from the spermatostyle was slightly broader than the other parts (Fig. 2T).

##### TRIBE LICININI

(Fig. 2U–X)

The formation of long spermatostyles was observed (Fig. 2U–W). Adhesion of the sperm was observed in a small part of the spermatostyle; the head of each sperm was attached to the spermatostyle, while the tail was free-moving (Fig. 2U). Other sperm were present that were not attached to the spermatostyle. The head of the spermatostyle was simple and did not broaden. The head of the single sperm did not broaden, and thus it was unclear which end was the head (Fig. 2X).

##### TRIBE OODINI

(Fig. 3A–C)

**Figure 3.**
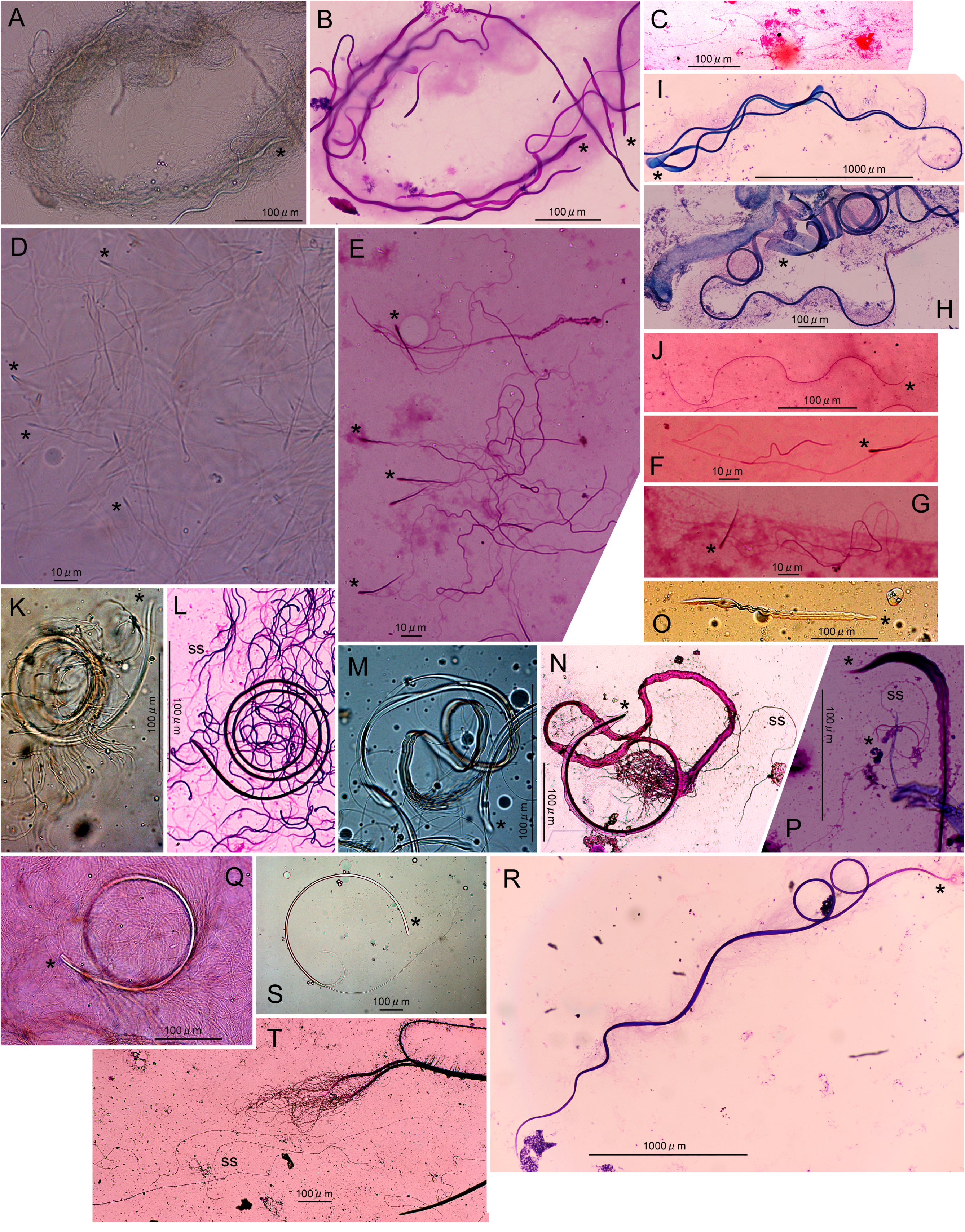
Sperm of various species of Carabidae: *Lachnocrepis prolixa* spermatostyles and single sperm in RS (A) and with GS (B), and single sperm with GS (C); *Agonum suavissimum* single sperm in RS (D) and with GS (E–G); *Xestagonum xestum* sperm conjugate with GS (H); *Metacolpodes buchanani* sperm conjugate (I) and single sperm (J) with GS; *Synuchus arcuaticollis* sperm conjugate in RS (K) and sperm conjugate and detached single sperm with GS (L); *Synuchus dulcigradus* sperm conjugate in RS (M) and sperm conjugate and detached single sperm with GS (N); *Synuchus orbicollis* sperm conjugate in RS (O) and sperm conjugate and single sperm with GS (P); an *Amara chalcites* sperm conjugate in RS (Q); an *Amara sinuaticollis* sperm conjugate with GS (R); *Amara macronota* sperm conjugate in RS (S) and sperm conjugate and detached single sperm with GS (T). Explanations of the markings are the same as in Figure 1.

Wave-shaped spermatostyles were observed; however, adhesion of the sperm to the spermatostyle was not observed (Fig. 3A, B). The head of the single sperm did not broaden, and thus it was unclear which end was the head (Fig. 3C).

##### TRIBE PLATYNINI

(Fig. 3D–J)

Two types of sperm were observed. The first type was observed in *Agonum suavissimum*, and only a single sperm was produced (Fig. 3D–G). The sperm had a long, drop-shaped head and two tails attached slightly behind the anterior end of the head. The tail was bifurcated in the rear half, and one of the bifurcated tails was broader than the rest of the tail and shorter than the other bifurcated tail (Fig. 3E–G). Motility was observed in Ringer’ s solution (Fig. 3D). The other type of sperm was observed in the other two species, and sperm conjugate was found. The spermatostyles had a long, left-handed helical shape and a spoon-shaped head (Fig. 3H, I). Numerous sperm attached their heads to the spermatostyle surface, while the tails were free-moving. The head of the single sperm detached from the spermatostyle was slightly broader than the other parts (Fig. 3J).

##### TRIBE SPHODRINI

(Fig. 3K–P)

Sperm conjugate, in which sperm were attached to spermatostyles, was observed, but the shape of the spermatostyles and the condition of the attached sperm differed between the species. In *Synuchus arcuaticollis* (Fig. 3K, L), the anterior and posterior halves of the spermatostyle had a left- and right-handed helical shape, respectively. Sperm were attached by their heads to the spermatostyle, while their tails were free-moving. The head of single sperm detached from the spermatostyle was distinctly broader than the other parts. In *Synuchus dulcigradus* (Fig. 3M, N), the posterior half of the spermatostyle had a left-handed helical shape, and the anterior half was oriented backward (compared to the helical direction of the posterior half). The sperm attached to the lateral sides of the spermatostyle along the midline and formed a sheet-like structure, except near the posterior end, where the tails of the sperm unraveled. The head of the detached single sperm did not broaden. *Synuchus orbicollis* formed a straight spermatostyle, and the sperm attached to the lateral sides of the spermatostyle, forming a sheet-like structure (Fig. 3O). The sheet-like structures undulated actively in Ringer’s solution. The heads of the single sperm detached from the spermatostyle were broadened (Fig. 3P).

##### TRIBE ZABRINI

(Fig. 3Q–T)

Sperm conjugate, in which sperm were attached to the spermatostyles, was observed; however, the shape of the spermatostyles and condition of the attached sperm differed between the species. In *Amara chalcites* (Fig. 3Q) and *Amara sinuaticollis* (Fig. 3R), the spermatostyles had a left-handed helical shape, and numerous sperm were attached by their heads to the spermatostyle surface, while the tails were free-moving. In *Amara macronota*, the anterior half of the spermatostyle formed a rigid and circular arc, while the posterior half was markedly slender and flexible (Fig. 3S). The sperm adhered to both of the lateral sides of the posterior half of the arc portion of the spermatostyle, while their tails were free-moving (Fig. 3T). The heads of the sperm that dissociated from the spermatostyle did not broaden in any of the three species.

##### SPERM SIZE VARIATION

Single sperm from 18 species, sperm conjugate from 15 species, and spermatostyles without attached sperm from 3 species were analyzed (Table 1). The size distribution of the sperm conjugate in *E. punctatus* deviated from the normal distribution (*D* = 0.106, *P* = 0.034), whereas the size distributions of all others did not deviate from normality (*P* > 0.05; Appendix S1). Comparison of the histograms and fitted smooth curves of the various sperm conjugate sizes between *E. punctatus* and the related species, *Elaphrus sugai*, revealed that the *E. punctatus* sperm conjugate had a positively skewed size distribution (Fig. 4).

**Figure 4.**
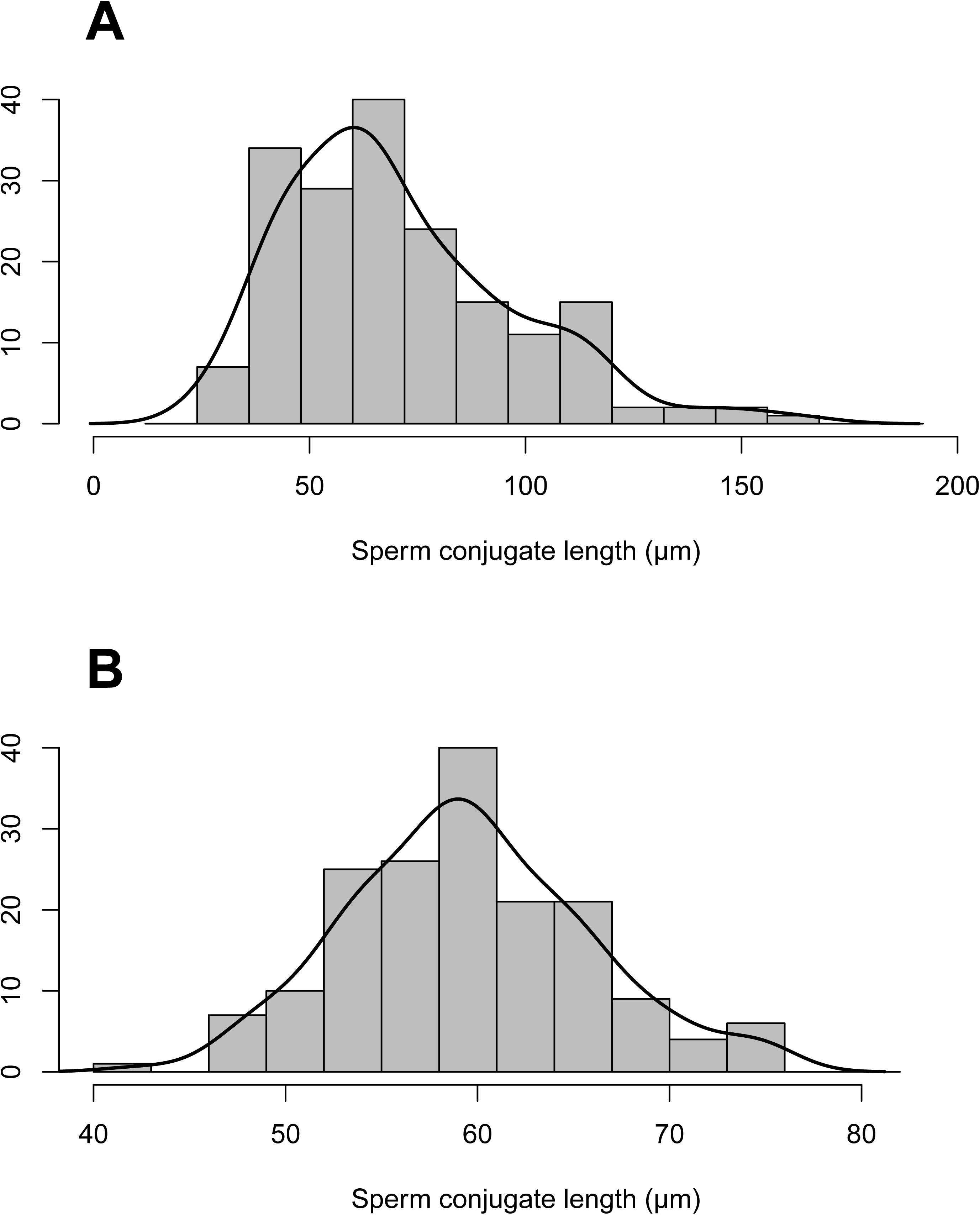
Variation in sperm conjugate length in *E. punctatus* (A) and *E. sugai* (B).

## DISCUSSION

Several notable sperm traits were uncovered in this study. First, the average length of the single sperm of *E. punctatus* was 55.10 μm. Prior to this study, the shortest single sperm reported in the order Coleoptera was that of the carabid beetle *Carabus maiyasanus*, which has an average length of 67.33 μm (Takami & Sota, 2007; Pitnick *et al*., 2009). Thus, the *E. punctatus* single sperm is now the smallest single sperm reported in Coleoptera. Furthermore, *E. punctatus* single sperm is also the second shortest in the class Insecta (Pitnick *et al*., 2009). The size variation in *E. punctatus* sperm conjugate is also notable. To date, size variation in sperm conjugate has been reported in various animal species, but only the sperm conjugate of carabid beetles of the genus *Carabus*, subgenus *Ohomopterus* has been quantitatively and statistically evaluated (Takami, 2002; Takami & Sota, 2007, both reported as “sperm bundles”). In *Ohomopterus*, the sperm conjugate exhibited species-specific variation in size dimorphism, ranging from a distinct bimodality to a positively skewed distribution (a lesser degree of bimodality); regardless of the degree of dimorphism, the sperm conjugate could be divided into two types: large and small (Takami & Sota, 2007). The positively skewed size distribution of the *E. punctatus* sperm conjugate may reflect a low degree of dimorphism; if so, this is the first report of dimorphism in the size of sperm conjugate, other than in *Ohomopterus*.

In the Harpalinae tribes Chlaeniini (except for one species), Dryptini, Galeritini, Licinini, and Oodini, few to none of the sperm were attached to the spermatostyles. One possible explanation for this observation is that, while loss of conjugation occurred in the species, the process of spermatostyle formation was maintained. If so, this sperm condition may represent a new type of sperm that evolved from the normal type of sperm conjugate. Another possible interpretation is that dissociation of the sperm from the spermatostyle is an artifact of the experimental procedures that were used. In this study, the sperm was surgically removed from the male seminal vesicles and observed in Ringer’s solution, which is the usual procedure for observing insect sperm (e.g., Hayashi & Kamimura, 2002; Higginson *et al*., 2012; Hodgson *et al*., 2013; Schubert *et al*., 2017). However, when sperm are transferred to the female, they are combined with other ejaculates, such as spermatophores. Physiological and/or chemical effects resulting from this difference may have caused the dissociation of sperm from the spermatostyles. Future studies should be conducted to examine the sperm in ejaculate collected from females immediately after mating in these species, as has been performed in other species (Takami, 2002; Takami & Sota, 2007; Sasakawa, 2007).

In the Harpalinae tribe Platynini, multiflagellated sperm were found, which have previously only been reported in one species in the class Insecta (Pitnick *et al*., 2009). The multiflagellated sperm previously reported were found in the Australian termite *Mastotermes darwiniensis* and had a conical head and about 100 simple (i.e., not branched) flagella (Baccetti & Dallai, 1978). Thus, the biflagellate sperm of *A. suavissimum* represent a second example of multiflagellated sperm in insects. The Platynini, which includes *A. suavissimum*, could serve as a model system for elucidating the evolutionary relationship between multiflagellated sperm and sperm conjugate, which may be mutually exclusive in terms of increased sperm motility; this is because aside from *A. suavissimum*, all Platynini species examined to date produced sperm conjugate consisting of uniflagellate sperm (Sasakawa & Toki, 2008; Schubert et al., 2017; Sasakawa, this study). Studies of sperm traits from additional Platynini species may provide further insights into this phenomenon.

In the coleopteran family Dytiscidae, which is the only example (to my knowledge) in which the evolution of sperm conjugation has been examined from a phylogenetic perspective, the same type of sperm conjugate as that observed in Carabidae was found to be the ancestral sperm trait, and other types of sperm conjugate evolved from it, followed by the loss of sperm conjugation in several lineages (Higginson *et al*., 2012). Compared to the findings for Dytiscidae, the current information regarding Carabidae sperm is insufficient, in that some phylogenetic relationships within the family are uncertain and the number of species examined for sperm traits is smaller. However, the available information revealed some of the evolutionary patterns of sperm traits in Carabidae. For the phylogenetic relationships, the latest subfamily-level phylogenetic trees based on mitochondrial genomes and nuclear 18S rRNA genes, which attempt to minimize the effects of various methodological problems (López-López & Vogler, 2017), and some robust sister group relationships that were confirmed in other molecular phylogenetic studies (Trechinae + Patrobinae: Maddison *et al*., 1999; Ribera *et al*., 2005; Brachinae + Harpalinae: Ribera *et al*., 2005; Ober & Maddison, 2008) are now available. These show that the relationship of taxa with known sperm traits is (Cicindelinae, (Carabidae, (Nebriinae, (Elaphrinae, ((Trechinae, Patrobinae), (Broscinae, (Scaritinae, (Brachinae, Harpalinae)))))))). Importantly, taxa with single sperm (i.e., Cicindelinae, Nebriinae, and Trechinae), and those with sperm conjugate (i.e., Carabidae, Elaphrinae, and Patrobinae), are placed alternately in basal lineages of the phylogeny. This implies that whether the ancestor of Carabidae formed single sperm or sperm conjugate is, at present, indeterminable; however, in either case, the transition between single sperm and sperm conjugate occurred several times in basal lineages. On the other hand, in Harpalinae, the largest subfamily that includes about half of the species of Carabidae, sperm conjugate with a long spermatostyle was considered most likely to be the ancestral trait, because most examined species of the subfamily and a species of Brachinae, the subfamily sister to Harpalinae, form sperm conjugate with a long spermatostyle.

The observed putative evolutionary pattern provides a definitive result for the phylogenetic utility of sperm morphology. Because two clades composed of species producing sperm conjugate with a long spermatostyle are separated in the phylogeny (i.e., Patrobinae in basal lineages, and Bachinae + Harpalinae at the most derived position), and an unambiguous transition from conjugated to single sperm has occurred in one of the two clades (*A. suavissimum* in the Harpalinae tribe Platynini), the current findings do not support Sasakawa & Toki’s (2008) assumption that a long spermatostyle contains phylogenetic information as an apomorphy. Also, sperm conjugate and single sperm are placed alternately on the basal lineages; thus, the macromorphology of sperm does not appear to contain useful phylogenetic information in Carabidae. To uncover phylogenetic information from sperm morphology, it will be necessary to examine micromorphology, such as the internal structure of sperm.

## ACKNOWLEDGEMENTS

I thank Drs Kôhei Kubota, Masahiko Tanahashi, and Wataru Toki for sample collections. The present study was partly supported by a grant-in-aid from the Japan Society for the Promotion of Science (nos. 20-11227, 25830150, and 17K15171).

